# The chloroplast located HKT transporter plays an important role in sporophyte development in *Physcomitrium patens*

**DOI:** 10.1101/2024.08.22.609226

**Authors:** Carolina Yanez-Dominguez, Karla Macedo-Osorio, Daniel Lagunas-Gomez, Diana Torres-CIfuentes, Juan Castillo-Gonzalez, Guadalupe Zavala, Omar Pantoja

## Abstract

Cell survival depends on the maintenance of cell homeostasis that involves all the biochemical, genomic and transport processes that take place in all the organelles within an eukaryote cell. In particular, ion homeostasis is required to regulate the membrane potential and solute transport across all membranes, any alteration in these parameters will reflect in the malfunctioning of any organelle. In plant cells, sodium transporters play a central role in keeping the concentrations of this cation across all membranes under physiological conditions to prevent its toxic effects. HKT transporters are a family of membrane proteins exclusively present in plants, with some homologs being present in prokaryotes. HKT transporters have been associated to salt tolerance in plants, retrieving any leak of the cation into the xylem, or removing it from aerial parts to be transported to the roots along the phloem. This function has been assigned as most of the HKT transporters are located at the plasma membrane. Here, we report the localization of the moss HKT from *Physcomitrium patens* to the thylakoid membrane, and its mutation that leads to several alterations in the phenotype of the organism, together with the changes in expression of close to 1000 genes. Down regulation of photosynthesis related genes and the upregulation of glycolysis/respiration and ion transport genes help to explain the observed phenotype.

## INTRODUCTION

A characteristic of all living cells is the presence of a membrane potential that is maintained by the interplay of transport proteins embedded in the plasma membrane, where primary ion transporters or ion pumps generate an electrochemical gradient across the membrane that energizes both, secondary active and passive transporters. While this is the case for membranes like the plasma membrane, tonoplast and that from organelles like endoplasmic reticulum and Golgi apparatus, in mitochondria and chloroplast, the electrochemical potential at the inner membrane of the former and the thylakoid membrane of the latter, is established by the electron transport chain activated by respiration or light, respectively. The thylakoid membrane, like any other biological membrane requires that all membrane transporters inserted within it work in a concerted manner to maintain an active light-driven electron transport chain during photosynthesis to sustain the synthesis of ATP and NADPH. Disruption of the electrochemical potential or proton motive force (pmf) would result in the inhibition of ATP and NADPH synthesis and therefore, photosynthesis.

As ATP synthesis directly depends on the ΔpH component of the proton motive force in the thylakoid, the electric component, ΔΨ, must be maintained at a required level to ensure a stable pmf to sustain ATP synthesis, as well as all the other processes that are activated during photosynthesis, like photo and oxidative protection. Several transport mechanisms have been proposed to serve for the dissipation of ΔΨ like the CLCe Cl transporter (Herdean et al., 2016), MSL ion channels (Haswell & Meyerowitz, 2006) or the KEA3 exchanger (Aranda Sicilia et al., 2021; Aranda-Sicilia et al., 2012; Kunz et al., 2014a; Uflewski et al., 2024). Presence of more than one of these transporters in the thylakoid, provides versatility for the adaption of the chloroplast to the continuous variations in environmental conditions.

The HKT family of plant-restricted ion transporters has been widely studied and described as mechanisms participating in salt tolerance as most of them function as low affinity sodium transporters located at the plasma membrane of xylem or phloem parenchymal cells, preventing, or removing sodium from the aerial parts of the plant (Horie et al., 2001, 2009). This family of transporters presents a fourfold membrane-pore-membrane (MPM) structure, grouped in two subfamilies, according to the presence of a S or G at the first pore of the selective filter, with the other three pores presenting a conserved G (Kato et al., 2001; Platten et al., 2006). Subfamily one, HKT1, possesses an S, while subfamily two, HKT2, presents a G at the first pore, functioning as Na^+^ transporters or Na^+^/K^+^ co-transporters, respectively (Kato et al., 2001; Platten et al., 2006). According to phylogenetic analyses (Riedelsberger et al., 2021), HKT’s are present from streptophyte algae up to angiosperms, grouped in three clades with sub-families1-2 separated from those transporters present in bryophytes (Riedelsberger et al., 2021). HKT’s from bryophytes present four G in the pore region of the selective filter, indicating that this structure is the ancestral version of the family, similar to the structure of the related Trk/Ktr prokaryote K^+^-selective transporters (Cao et al., 2011, 2013). In agreement to the K^+^-selectivity associated with the presence of four G in the pore regions, the bryophyte HKT homologues characterised so far, transport both K^+^ and Na^+^ (Haro et al., 2010a; Imran et al., 2022), however, the physiological or morphological role for these transporters is not clear. Here, we describe the morphological changes caused by mutation of the single *Pp*HKT present in *Physcomitrium patens*, as well as its localization to the chloroplast, together with an RNAseq analysis that indicate the mutation of this gene has important consequence in photosynthesis and several other mechanisms that help to explain the pleiotropic changes observed in the *PpΔhkt* mutant.

## RESULTS

### Mutation of *PpHKT* generates pleiotropic effects at all stages of the moss life cycle

Transporters from the HKT family have been described as K^+^ or Na^+^ mechanisms located at the plasma membrane, with some exceptions, involved in the absorption of these two cations and associated with high affinity K^+^ and/or low affinity Na^+^ uptake. In a previous study, it was reported that *Pp*HKT does not seem to play an important role in K^+^ or Na^+^ uptake in the moss *P. patens*, as growth was unaffected by the mutation of the corresponding gene. In that report, however, there was no description on morphological changes caused by the mutation, in particular, at later stages of its life cycle, which led us to analyze the morphology of the mutant. Moss development from the *PpΔhkt* line at 7 days after protoplast regeneration looked bigger and with more branches than the WT (Fig. S1A); after 10 d, these differences were more obvious (Fig. S1A), and were confirmed by quantifying the surface area covered by the protonema (Fig. S1B). When the moss was grown in colony and under long day conditions, gametophores from the *PpΔhkt* were almost twice as big as those from the WT line (compare Fig. 1A-B; left panels and Fig. 1C); while the difference in size between gametophores grown on short day conditions from the two lines was not as large, but still observing a smaller size in those formed by the WT line (compare Fig. 1A-B; right panels and Fig. 1C). Additional differences were observed for the gametophores from *PpΔhkt* mutant, which presented a thinner stem and smaller phyllids with a more vertical position with respect to the stem (Fig. 1A-B, right panel). The most conspicuous difference between the two lines was the absence of sporophyte formation in the mutant line (Fig. 1A-B, right panels). Confirmation of these differences was obtained by quantifying gametophore length and percent of gametophores with sporophyte (Fig. 1C-D). The almost complete absence of sporophytes in the *PpΔhkt* mutant is shown in Figure 1D, while observing that the WT line generated sporophytes in almost 50% of the gametophores, a value in agreement with previous reports (Daku et al., 2016; Singer & Ashton, 2007). Finally, gametophores from the *PpΔhkt* mutant grew under short day conditions, were paler in color in comparison to the WT (Fig. 1A-B, right panels), because of a lower chlorophyll content (Fig. 1E).

**Figure 1.**
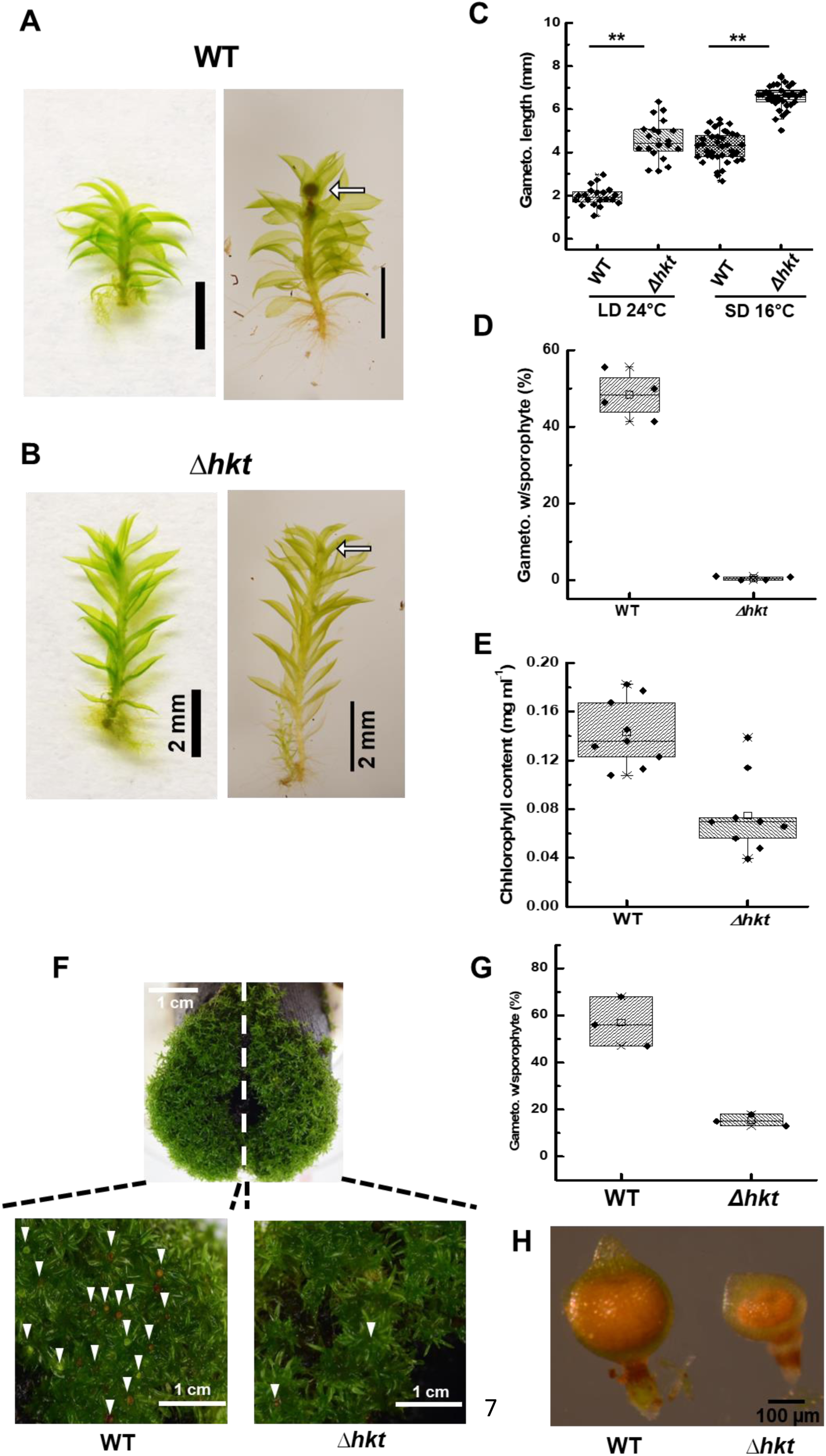
Deletion of *PpHKT* generates pleiotropic changes at the gametophyte and sporophyte stages. **A** Wild type (WT) gametophores grew on PpNH_4_formed normal adult gametophores after 30 d under a long day photoperiod (LD) condition (left panel) and WT gametophores growth on Jiffy pellets after sporophyte induction under a short-day photoperiod (SD) condition shows the development of normal sporophyte (right panel, arrow). **B** The *PpΔhkt* gametophores grew on PpNH4 medium formed larger gametophores after 30 d under a LD condition (left panel), and *PpΔhkt* gametophores grew on Jiffy pellets after sporophyte induction under SD conditions did not produce sporophyte (right panel, arrow) **C** *PpΔhkt* gametophores (n=34) were longer than WT (n=38), independently of the growth conditions. t-test, p-value < 0.001. LD, SD. **D** Fifty percent of WT gametophores (n= 412, data from four independent experiments) produced a sporophyte, while those from the *PpΔhkt* mutant (n= 474, data from five independent experiments) did not produce any sporophyte. t-test p-value< 0.001. **E** Gametophores from the *PpΔhkt* line (n=30) contain less chlorophyll than the wild type. Data from nine independent experiments, bars indicate standard deviation: t-test p-value< 0.05. **F** Co-cultivation of WT and *PpΔhkt* lines caused the formation of sporophytes in both lines (arrows). **G** Close to 20% of the gametophores from the *PpΔhkt* mutant produced sporophytes, while in the WT, 60% did **H** Sporophytes from the *PpΔhkt* line were smaller and deformed, in comparison to WT.

To identify if the lack of sporophyte formation in the mutant *PpΔhkt1* was due to a malformation of the sexual organs, we analyzed their development in both lines. From 15 days post-induction (dpi), up to 28 dpi, the archegonia and antheridia from both lines showed similar development (Fig. S2), indicating that the absence of sporophyte formation in the *PpΔhkt* mutant may be related to the fertilization stage. To confirm or discard this possibility, the WT and mutant lines were grown in the same Jiffy pellet, expecting to induce the formation of sporophytes in the mutant by fertilization with the WT spermatozoid. Under these conditions, abundance of sporophytes was normal in the WT moss (Fig. 1F, left; Fig. 1G), while observing the presence of few sporophytes in the *PpΔhkt* mutant (Fig. 1F, right), that accounted for an average of 15% of gametophores with sporophytes (Fig. 1G). Interestingly, the sporophytes formed in the *PpΔhkt* mutant were smaller than those from the WT line (Fig. 1H). These results indicate that *Pp*HKT might be important for fertilization and the development of the sporophyte.

### *Pp*HKT locates to the chloroplast

After observing these strong modifications in the phenotype of the *PpΔhkt* mutant, we identified the intracellular localization of the transporter by generating the *Pp*HKT-3xmNeon knock-in line. As shown in figure 2, the fluorescence associated with the transporter was observed as puncta within the chloroplast of protonemata cells, indicating its possible association with the thylakoid (Fig. 2A). The localization of *Pp*HKT-3XmNeon in the phyllids was patchy, with some areas showing groups of cells where the fluorescence was clearly associated with the chloroplasts, either, throughout the lamina (Fig. 2B, top) or at the edge of the phyllid (Fig. 2B, bottom). It is important to mention that chloroplasts that showed fluorescence associated with *Pp*HKT3XNeon did not show chlorophyll autofluorescence and had a different shape than chloroplasts present in the surrounding cells (Fig. 2B). Additionally, these differences in the fluorescence and shape of the chloroplasts, coincided with their brightness in the brightfield image, where the fluorescence of *Pp*HKT3XNeon was associated with brighter chloroplasts (Fig. 2B). Additional information on the localization of *Pp*HKT3XNeon was obtained by observing its presence in the reproductive organs, the archegonia, and the antheridia (Fig. 2C, D). In archegonia, fluorescent puncta were visualized along the neck cells, sometimes associated to chloroplast (Fig. 2C, square) or not (Fig. 2C, arrows), indicating the possible association of *Pp*HKT-3xmNeon with plastids. In antheridia, *Pp*HKT-3xmNeon fluorescence was observed in the spermatids that form between stages 7 and 8 inside the cavity, according to (Landberg et al., 2013). The spermatids were clearly identified by the presence of a central large nucleus (Fig. 2D, DAPI). *Pp*HKT3XNeon fluorescence was also observed inside the outer cell layer associated with small vesicles (Fig. S3). These results demonstrate that *Pp*HKT is located in the chloroplast, very likely in the thylakoid membrane, according to the intraplastid fluorescence signal, but also located in plastids, as suggested by the presence of *Pp*HKT inside non-fluorescent organelles.

**Figure 2.**
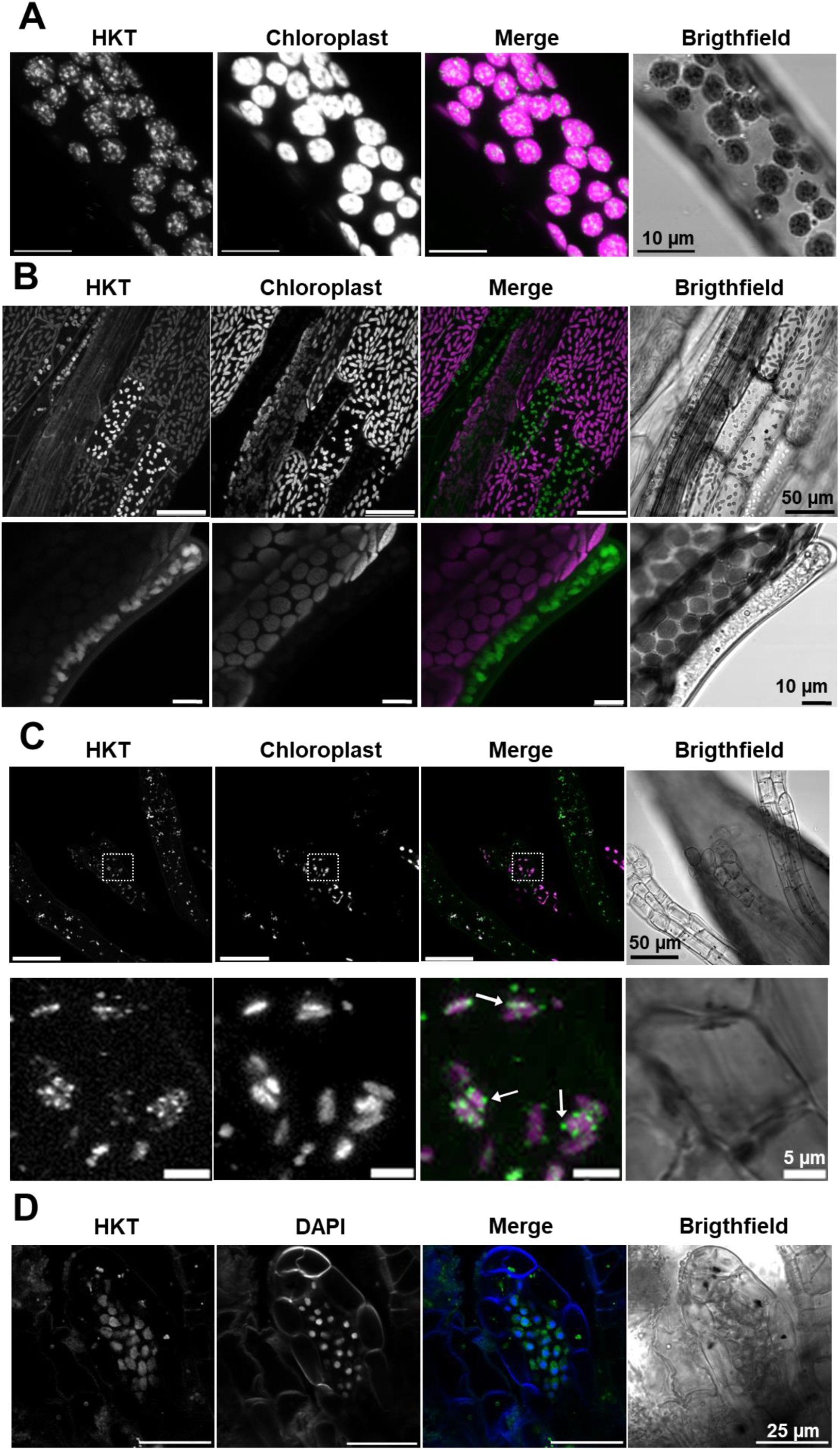
*Pp*HKT resides at the chloroplast. **A)** In protonema cells, HKT-3xmNeon (HKT) localizes in dots inside the chloroplast (Chloroplast), as shown in the merged images (Merge). **B)** In phyllids, HKT-Neon is observed in cells at the centre (top) or edge (bottom) of the lamina. **C) I**n arquegonia, HKT-3xmNeon localizes at plastids in cells of the neck (top); the square areas are shown as enlarge images (bottom). **D)** In antheridia, HKT-3xmNeon localizes in the sperm cells (HKT), as indicated by staining of the nuclei (DAPI) and confirmed in the merged images. Images in **A**, **B** and **C** correspond to Z-stack images showing HKT in green and chloroplasts in magenta. Images in **D** are stacks of 10 confocal slices from the central part of the antheridium.

### Mutation of *PpHKT* modifies chloroplast morphology

Localization of *Pp*HKT at the thylakoid membrane, together with a lower chlorophyll content and paler color of the gametophores from the *PpΔhkt* mutant, suggested malfunctioning of this organelle as the mechanism responsible for all these alterations. To identify possible changes in the morphology of the chloroplast, we analyzed these organelles at the ultrastructural level by transmission electron microscopy (TEM). The morphology of chloroplast from the phyllids of the WT moss showed the typical structure characterized by the oval shape of the chloroplast containing an extensive network of photosynthetic membranes (thylakoids), parts of which are appressed into moderate granal stacks, as well as the presence of several starch grains (Fig. 3A, top). Another group of chloroplasts did not show well defined thylakoid membranes, which were only seen at higher magnification (Fig. 3A, bottom). In both types of chloroplasts, the presence of large starch grains was also a common feature. A contrasting morphology was identified for the chloroplasts from the *PpΔhkt* mutant as indicated by alterations in the structure of the thylakoid membrane, losing the parallel arrangement of the lamella and without a clear definition of the chloroplast envelop (Fig. 3B, top). Like in the WT, a different type of chloroplast was also observed, characterized by not showing clear thylakoids but with the presence of several plastoglobuli and the absence of starch granules (Fig. 3B, bottom). These results are clear evidence that absence of *Pp*HKT from the thylakoid membrane affects its structure and probably, chloroplast functioning.

**Figure 3.**
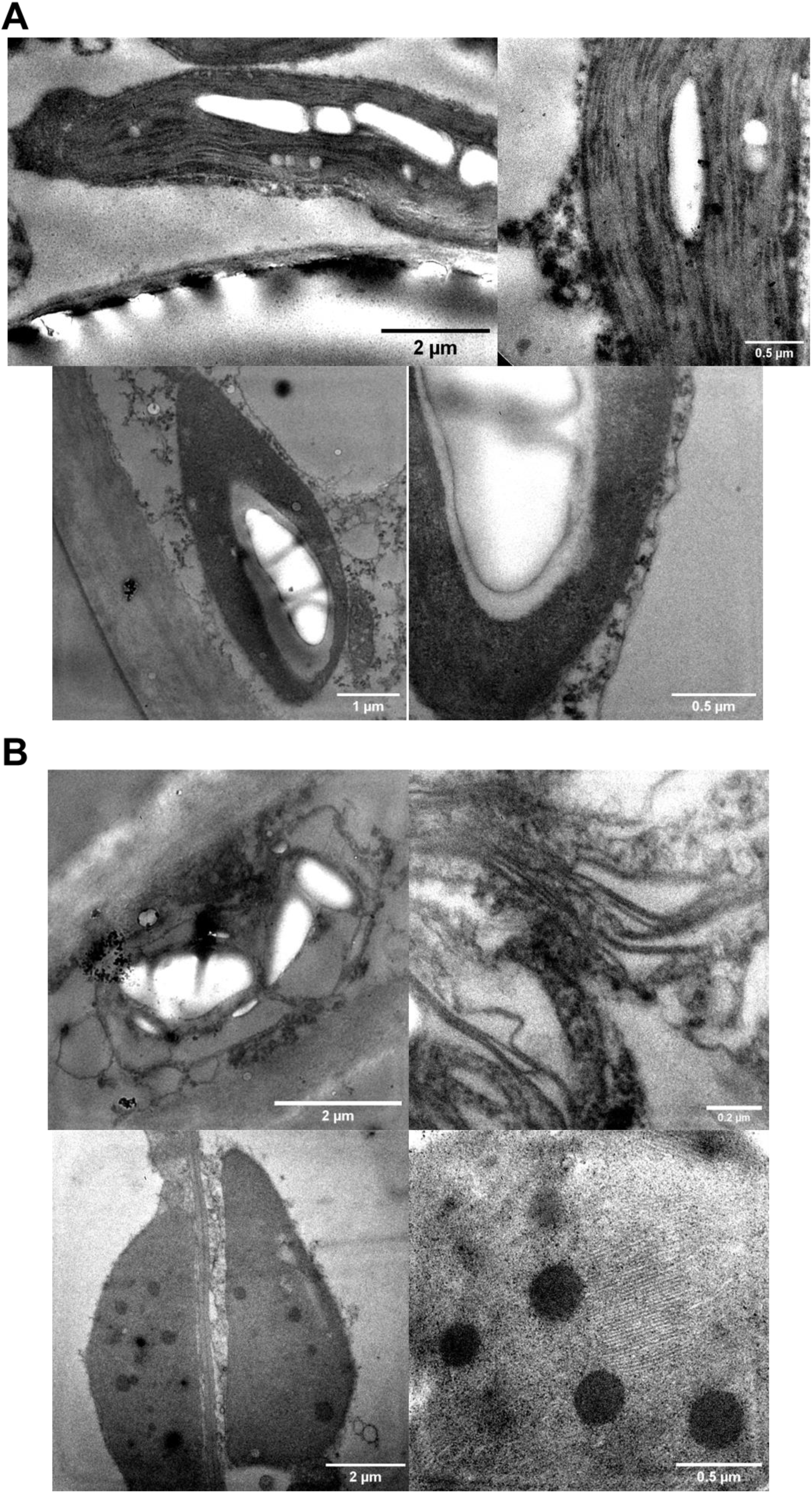
Chloroplast ultrastructure is modified in the *PpΔhkt* mutant. Chloroplast morphology from WT **A** and *PpΔhkt* **B** moss lines. Observe the unstructured thylakoid membrane and the presence of plastoglobuli in the *PpΔhkt* mutant.

### *Pp*HKT transport both, K^+^ and Na^+^

Previous characterization of *Pp*HKT has shown that it functions as a K^+^ or Na^+^ transporter (Haro et al., 2010b), a result we confirmed by its heterologous expression in yeast cells, after failing to record any activity from Xenopus oocytes injected with *Pp*HKT cRNA. Transforming the K^+^-uptake deficient yeast mutant BYT12 (*MATa his3Δleu2Δ0 met15Δ0 ura3Δ0; trk1::loxP trk2::loxP*) with *PpHKT* allowed the growth of the cells in YNB medium containing nominal K^+^ (Fig. 4A; 0 KCl). In comparison, BYT12 cells transformed with the empty vector required 20 mM K^+^ to show a clear growth (Fig. 4A). A similar approach was employed to evaluate the capacity of *Pp*HKT to transport Na^+^ employing the yeast BYT45 strain (*MATa his3Δ1 leu2Δ0 met15Δ0 ura3Δ0 nha1Δ::loxP ena1-5Δ::loxP*), whose growth is inhibited at high Na^+^ concentrations. In BYT45 cells, cell growth was clearly inhibited at 200 mM Na^+^ (Fig. 4B), an effect that was enhanced by the expression of *PpHKT*, where cell growth was repressed at 100 mM (Fig. 4B). The higher sensitivity of *PpHKT* transformed cells to Na^+^ was confirmed by the inhibition of cell growth at 20 mM Li^+^, in contrast with the cells transformed with the empty vector (Fig. 4C).

**Figure 4.**
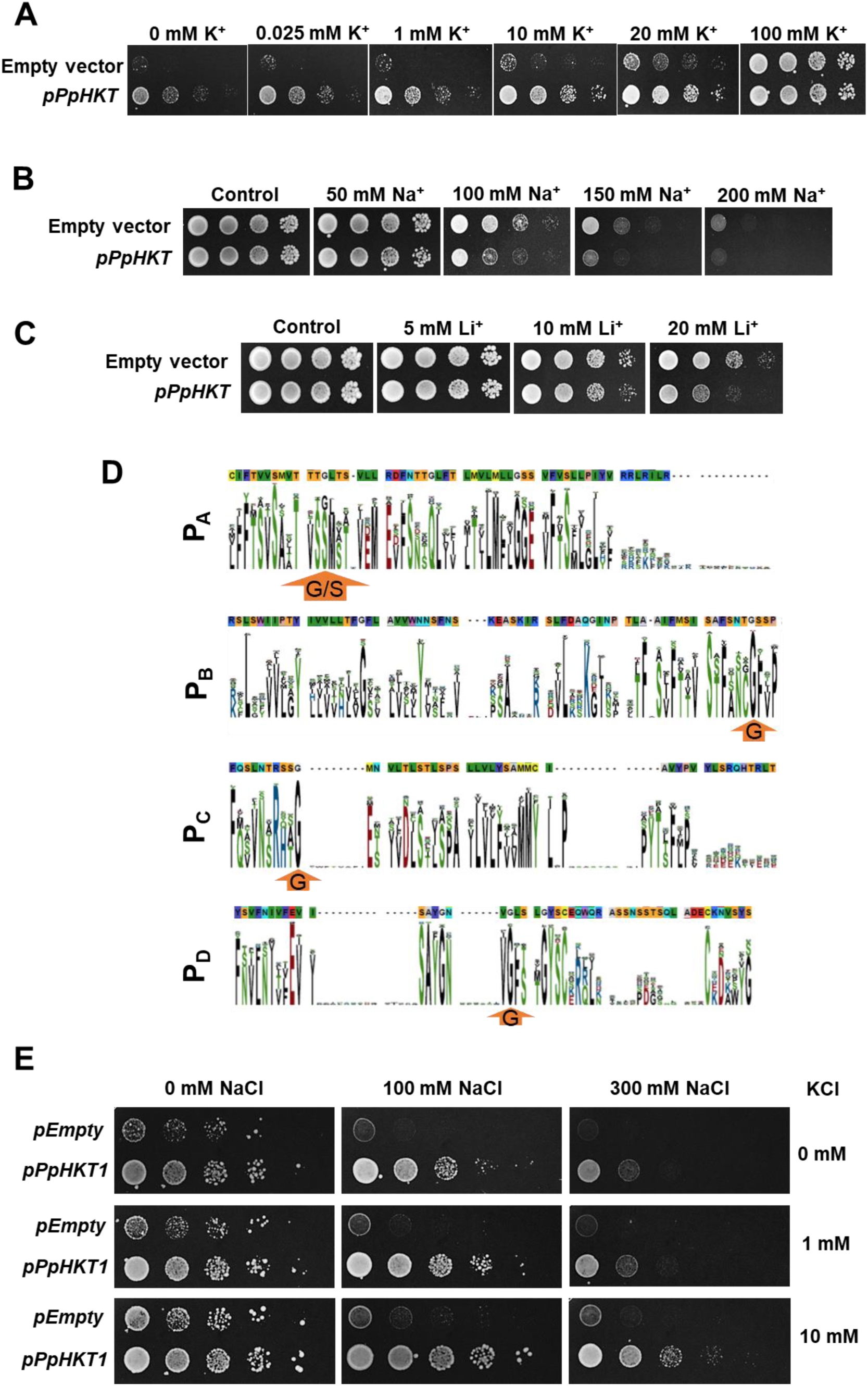
*Pp*HKT functions as a high affinity K^+^ transporter and as a low affinity Na^+^ transporter. **A** BYT12 yeast cells (*Δtrk1 Δtrk2*) did not grow at μM concentrations of K^+^ when transformed with the empty vector but grew when expressing *Pp*HKT. The sensitivity of the BYT45 yeast strain to Na^+^ **(B)** or Li^+^ **(C)** was increased by the expression of *PpHKT*. **(D)** Location of the pore located Gly residues in *Pp*HKT1 (top line) and their conservation in plant homologs (sequence logo). **(E)** Sodium sensitivity of the BYT45 yeast strain was decreased in the presence of increasing concentrations of K^+^.

According to protein sequence analysis (Fig. 4D), *Pp*HKT possesses a Gly in each of the four pore-forming domains, a property that has been associated with the function of homologue transporters as K^+^-Na^+^ co-transporters (Golldack et al., 2002; Horie et al., 2001). To test if *Pp*HKT was able to co-transport both cations, BYT12 cells were grown in varying concentrations of both cations, observing that the toxic effect of Na^+^ was diminished as the concentration of K^+^ was increased (Fig. 4E), indicating that both cations seemed to be transported by *Pp*HKT.

### The *PpΔhkt* mutant shows a modified transcriptome

To identify genes underlying the differences associated to the *PpΔhkt* phenotype, we performed a comparative RNAseq analysis from WT and *PpΔhkt* mutant lines. Mapping of sequenced samples resulted in alignment rates between 94–96% with v.3.3 of the *Physcomitrium patens* genome. When comparing gene expression between WT and *PpΔhkt* strains in a volcano plot employing a log_2_ (FoldChange) ≥±1 with a log p_adj_ ≤ 0.05, it was clearly observed the up- or down-regulation of 924 genes (Fig. S3A), of which 604 were down-regulated, while the expression of 320 genes was up-regulated (Fig. S3B). Cluster analysis from these differentially expressed genes allowed the identification of four clusters that were characterized by the abundance of genes related to ion transport, microtubule-based processes, photosynthesis, and cell projection organization (Fig. S3C). The genes associated with the first cluster, ion transport, were up-regulated, while those from the other three groups were down-regulated, with the largest number of down-regulated genes associated with photosynthesis (Fig. S3C).

Additional gene enrichment analysis was performed with the ShinyGO 0.8 algorithm (http://bioinformatics.sdstate.edu/go, (Xijin Ge et al., 2020)) employing an FDR cutoff value of 0.05 for identified pathways, which allowed us to obtain deeper information on gene alterations that could help to explain the phenotypes observed for the *PpΔhkt* mutant. We separately analyzed the down- and up-regulated genes. According to the number of downregulated genes and the -log (FDR) values for significant pathways derived from the ShinyGO analysis that classifies the DGE into three groups, Biological Process (BP), Molecular Function (MF) and Cellular Component (CC) (Fig. 5-7), most of the down-regulated genes in the *PpΔhkt* mutant corresponded to photosynthesis or photosynthesis related processes. In the BP group, the GO terms photosynthesis and generation of precursor metabolites comprise genes that code for photosystem I and II reaction center subunits, Rubisco small chain and oxygen-evolving enhancer proteins (Fig. 5B). The second most representative pathway within the BP group was Generation of precursor metabolites and energy that include pathways resulting in the formation of precursor metabolites and any process involved in the liberation of energy from these metabolites (Fig. 5C). Two additional GO-terms within the BP group were enriched, cell projection organization, that include genes coding for several dynein and dynein-related proteins, together with intraflagellar transport proteins (Fig. 5A, yellow arrow and Fig. 6E), and translation, mainly represented by ribosomal proteins, as well as others involved in protein folding (Fig. 5A, red arrow and Fig. 5D). In the MF group (Fig. 6A), chlorophyll binding and photosynthesis related genes from both, light and dark reactions, were down-regulated (Fig. 6A), together with genes involved in lipid metabolism and glycolysis comprised within the GO term oxidoreductase activity (Fig. 6B). Genes belonging to the GO term structural constituent of ribosome were down-regulated as well (Fig. 6A), in agreement with the attenuation of genes associated to translation identified within the BP group (see above). According to the down-regulated genes in the BP and MF groups, those within the CC group were mainly associated with the thylakoid or the chloroplast (Fig. 7A-B). The down-regulation of all these genes in the mutant line *PpΔhkt1* shows that the main process affected in the *PpΔhkt* mutant is photosynthesis, strengthening the localization of the transporter to the thylakoid membrane.

**Figure 5.**
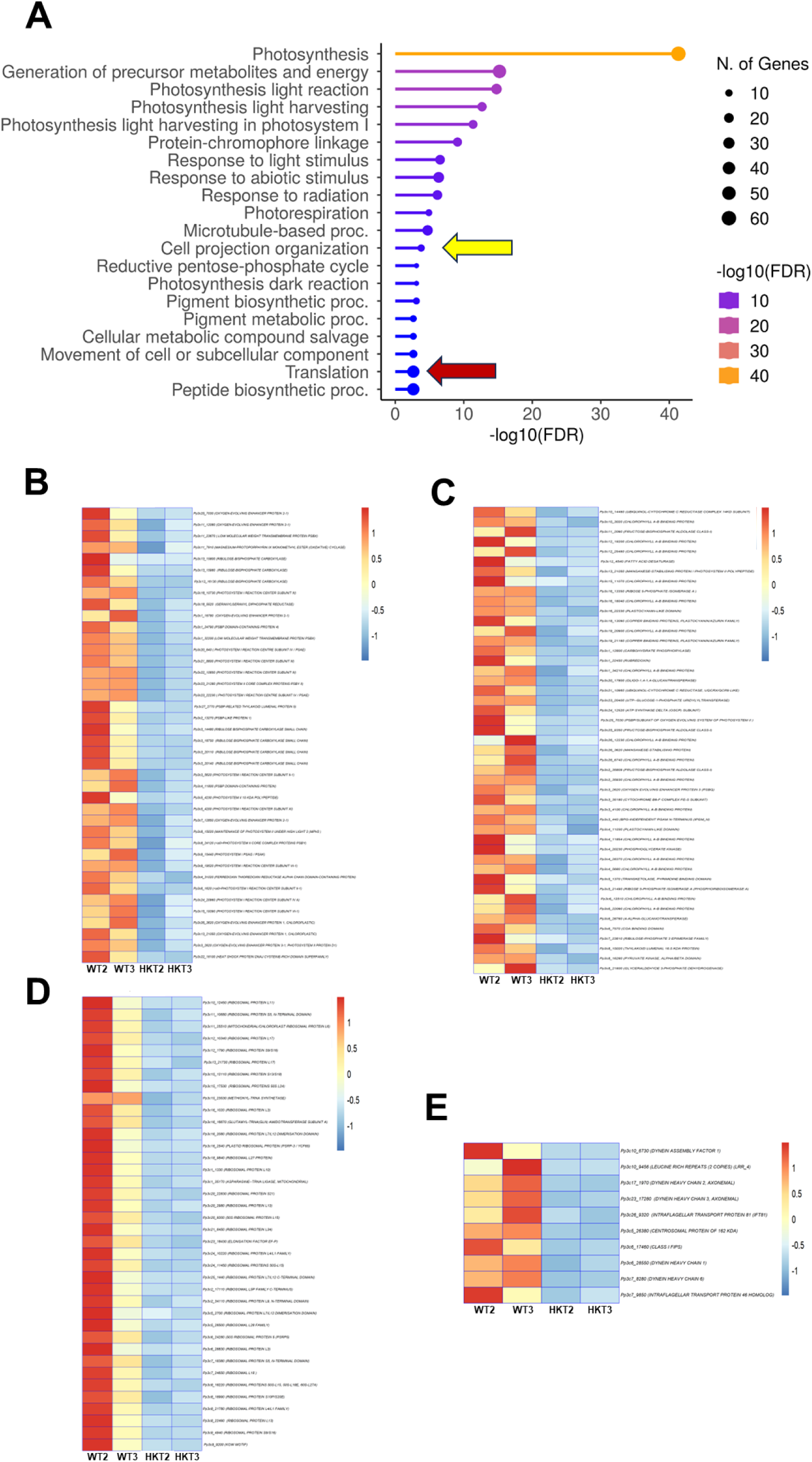
Dow-regulated Biological Processes in *PpΔhkt.* **A** Affected pathways were mainly associated to photosynthesis, but also including flagella (yellow arrows) or translation (red arrows). **B** Down-regulated genes related to Photosynthesis. **C** Down-regulated genes related to Generation of precursor metabolites and energy. **D** Down-regulated genes related to Translation in chloroplast or mitochondria. **E** Down-regulated genes related to Cell projection organization.

**Figure 6.**
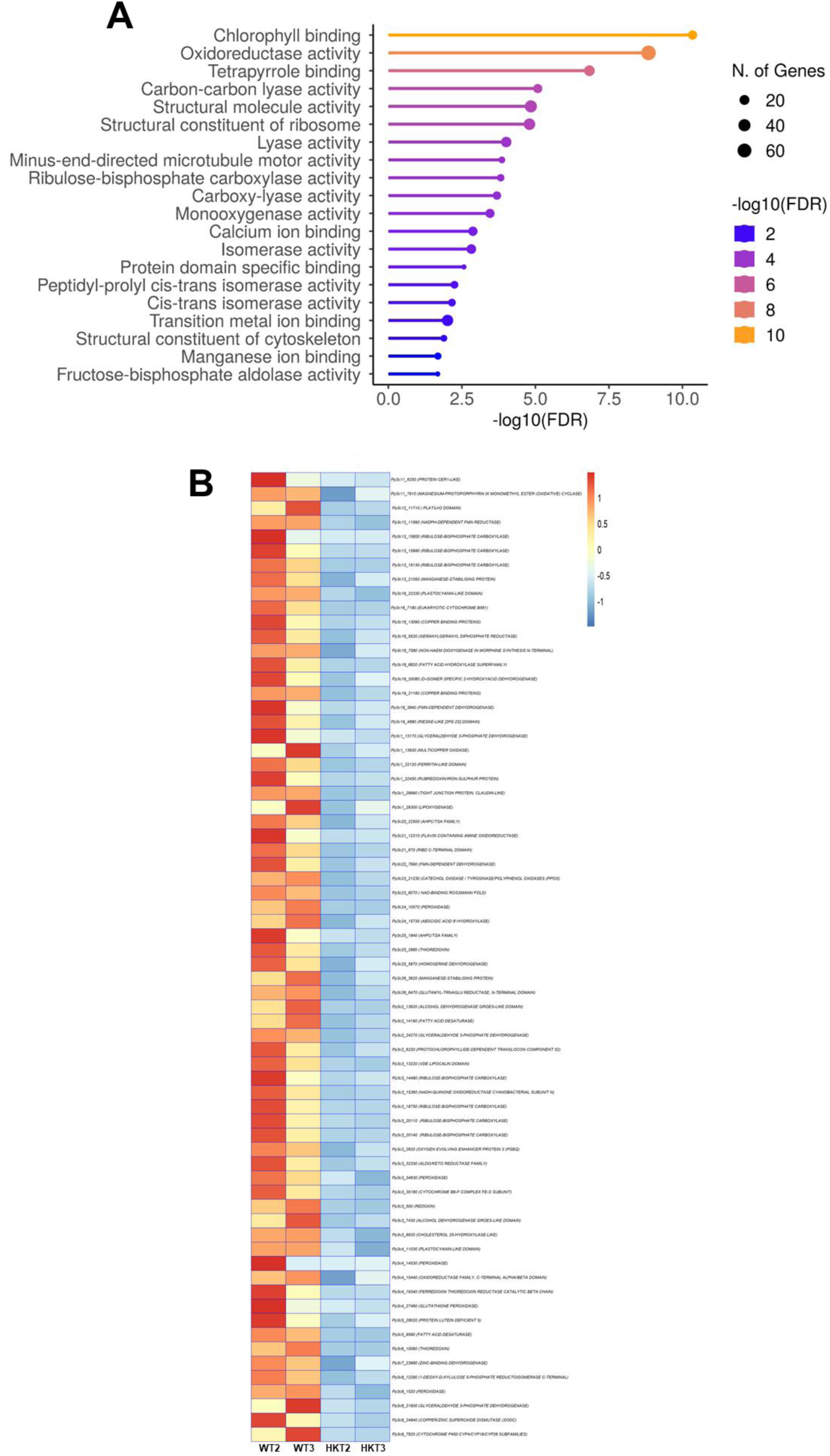
Down-regulated Molecular Functions in *PpΔhkt.* **A** Most of the molecular functions down-regulated in *PpΔhkt* were mainly associated to chlorophyll binding or oxidoreductase activities). **B** Down-regulated genes associated with Oxidoreductase activity.

**Figure 7.**
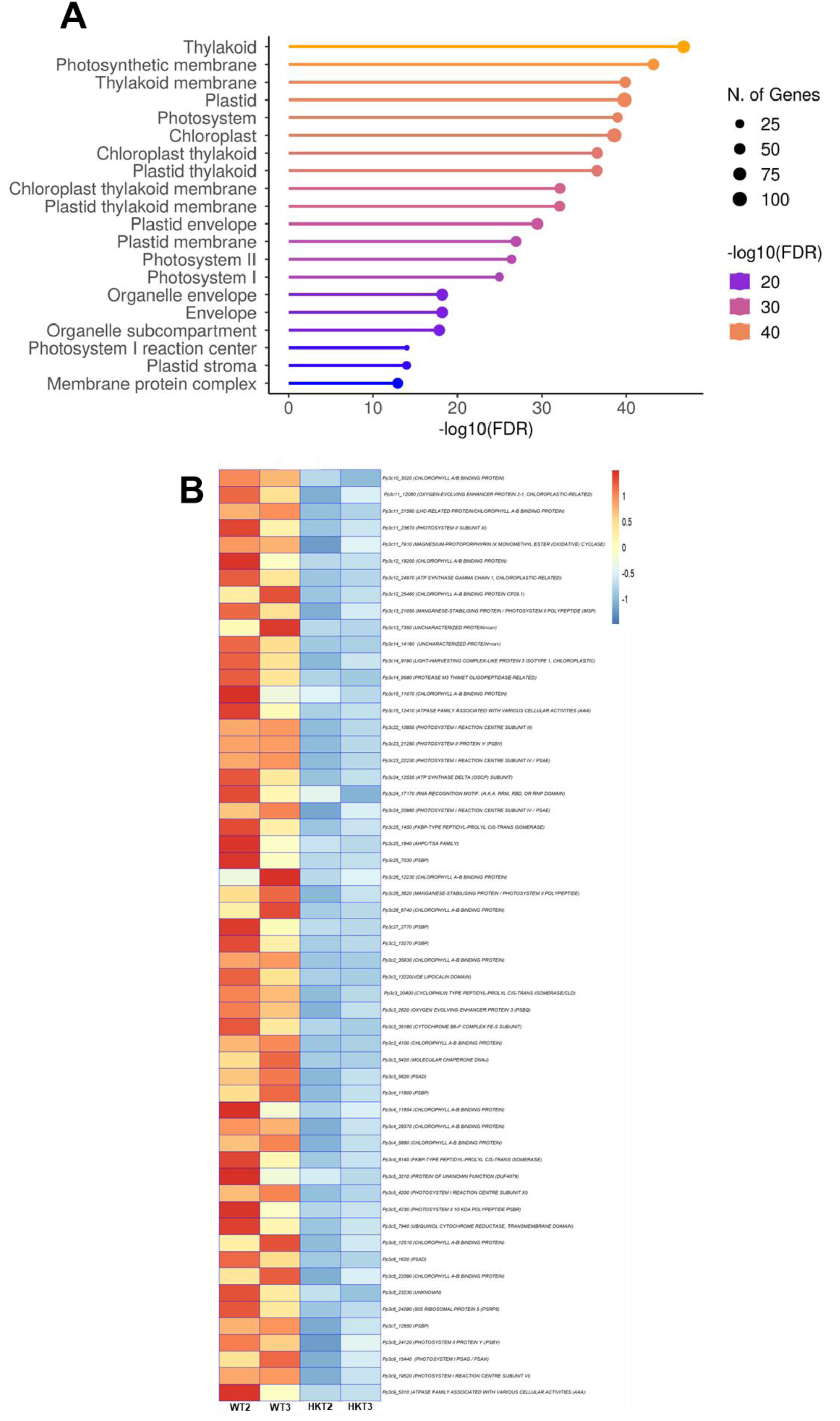
Down-regulated Cellular Components in *PpΔhkt.* **A** Most of the cellular components down-regulated in *PpΔhkt* were associated to the thylakoid membrane. **B** Down-regulated genes associated to the thylakoid.

Regarding the expression of upregulated genes, analysis of the GO terms clearly indicated that the main process modified by mutation of the chloroplast located *Pp*HKT transporter, was ion transport (Fig. 8), with particular enrichment in Zn^2+^ and K^+^ transporters within the Biological Process (Fig. 8A) and Molecular Function (Fig. S4A) groups. Several genes from the ABC Transporter and AAA+ ATPase families were also significantly upregulated (Fig. 8A-B). Also, within the BP group, three gene sub-groups, oxoacid, organic acid and carbohydrate metabolic processes, which are related to lipid, carbon and amino acid metabolism were upregulated in the *PpΔhkt* mutant (Fig. 8B-E). Finally, the expression of an important number of transcription factors was also increased, with half of them belonging to the AP2/ERF family (Fig. S4B). In *P. patens*, these transcription factors determine stem cell identity and cell reprogramming, acting as downstream factors by chromatin modification (Aoyama et al., 2012; Ishikawa et al., 2019; S. Wang et al., 2020).

**Figure 8.**
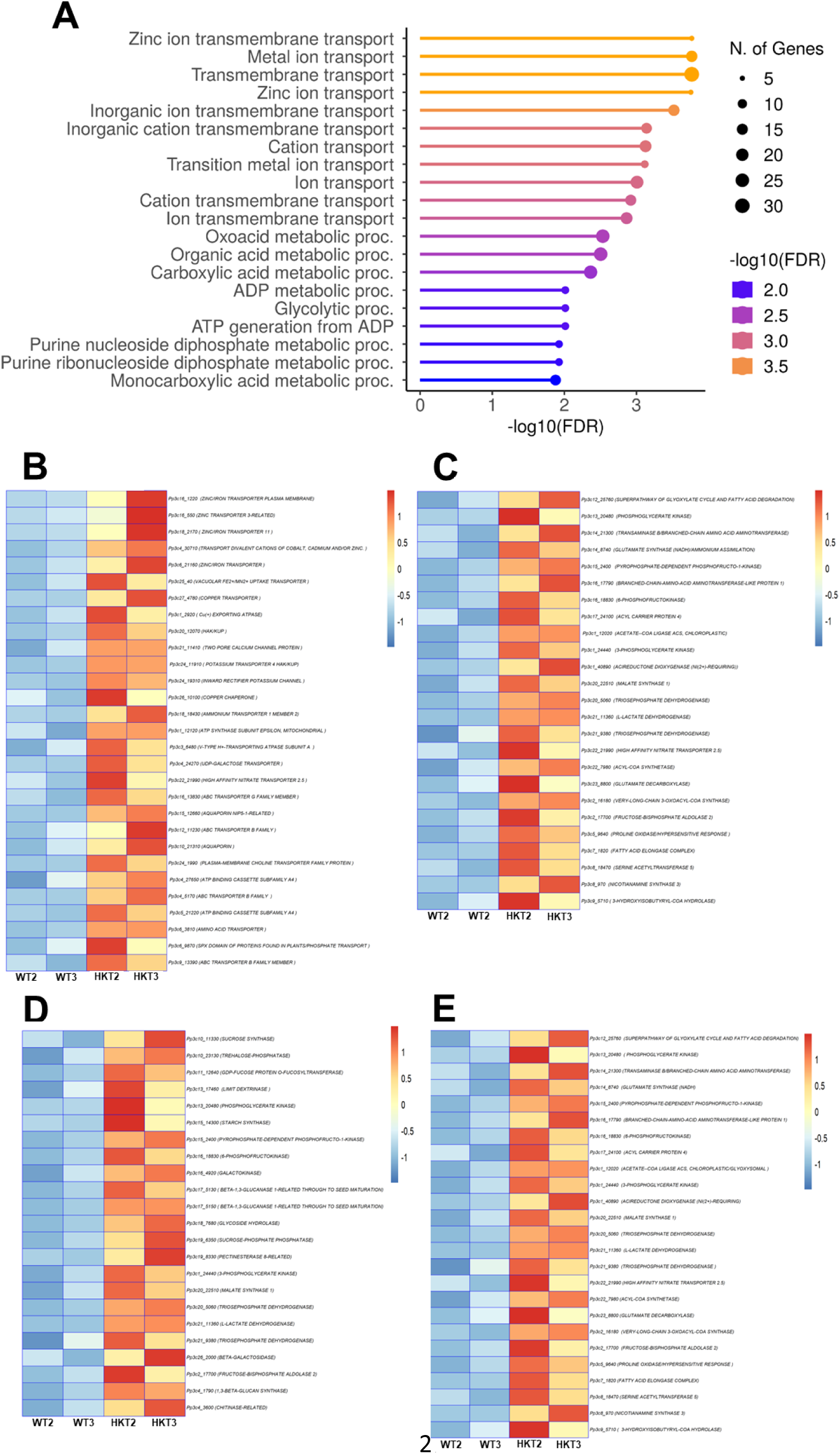
Up-regulated Biological Processes in *PpΔhkt.* **A** Most of the up-regulated biological processes in *PpΔhkt* were associated to **B** transmembrane transport, **C** Oxoacid metabolic process, **D** carbohydrate metabolic process or **E** organic acid metabolic processes.

## DISCUSION

Ion transport across biological membranes plays an important role in maintaining the ionic homeostasis within the organelles for the well-functioning of a cell. HKT transporters are a protein family present exclusively in plants, where it has been proposed to play a central role in salinity tolerance by preventing the long-distance transport of the toxic cation Na^+^ from the roots to the shoots, or in the high affinity uptake of K^+^, as they have been mainly localized to the plasma membrane. In contrast to these observations, we identified the localization of *Pp*HKT to the chloroplast, probably associated to the thylakoid membrane, according to the intra-chloroplast signal observed with the knock-in HKT-3XmNeon reporting line (Fig. 2). In agreement with the presence of four Gly in the pore domains, *Pp*HKT transports both K^+^ and Na^+^ with a higher affinity for the former (Fig. 4), and possibly, functions as a K^+^/Na^+^ co-transporter (Fig. 4), just as it has been reported for members of the sub-family HKT2 from monocots. Localization of *Pp*HKT to the thylakoid membrane, together with the morphological changes observed in the *PpΔhkt* mutant, clearly indicate a central role for this transporter in controlling ion homeostasis in the chloroplast, and therefore, in the physiology of the moss. Chlorosis and a lower chlorophyll content in the *PpΔhkt* mutant gametophore (Fig. 1) are the most obvious consequence of a malfunctioning chloroplast that results in additional alterations, like the absence in the formation of the sporophyte (Fig. 1) and the altered expression of close to 900 genes (Fig. 5). From these, most of the downregulated genes are associated to photosynthesis, with many genes related to chlorophyll synthesis, like magnesium-protoporphyrin IX monomethyl ester (oxidative) cyclase (Pp3c11_7910), which catalyses the formation of the isocyclic ring in chlorophyll biosynthesis; a magnesium protoporphyrin IX methyltransferase (ChlM, Pp3c12_24890), involved in the later steps of chlorophyll biosynthesis; a geranylgeranyl diphosphate reductase (Pp3c18_5520), involved in the reduction of geranylgeranyl diphosphate to phytyl diphosphate, providing a phytol molecule for Chlorophyll (Chl) synthesis. Many genes codifying for the light-harvesting chlorophyll complex LHCA and LHCB were also down-regulated in the *PpΔhkt* mutant (Fig. 6A-B), together with several oxygen-evolving-complexes (OEC) such as PsbP and PsbQ, and chloroplast electron transport components (Fig. 5-7), these changes could explain the chlorotic phenotype of the *PpΔhkt* mutant (Fig. 1). Moreover, according to the phenotype and the changes in gene expression observed for the *PpΔhkt* mutant, it is very likely that water photolysis together with a disruption of the electron flow in the thylakoid membrane result in a decrease in the reduction of NADP^+^ to NADPH and/or decreased ATP synthesis. This condition can generate reactive oxygen species (ROS) that can lead to the oxidation of a variety of cellular processes, molecules, that result in the pleiotropic morphological changes observed for the *PpΔhkt* mutant, where ROS could function as a retrograde signal (De Souza et al., 2017). In agreement with this reasoning, RNAseq results showed that a chloroplast superoxide dismutase (Fe-SOD) was the most upregulated gene (Pp3c17_14510), together with a cooper chaperone for superoxide dismutase (Pp3c17_14500), with a ten- and seven-fold increase, respectively, as mechanisms to ward off from the ROS.

A decrease of energy molecules like NADPH and ATP, because of a broken photosynthesis would lead to a likely disfunction of the Calvin cycle for carbohydrate synthesis. This was confirmed by the downregulation of several genes related to the Calvin cycle, like several isoforms of the small subunit of RuBisCO (Fig. 5-7). Confirmation that chloroplast functioning is widely affected in the *PpΔhkt* mutant is the decreased expression of genes related to protein synthesis in the chloroplast, in particular, but also in the mitochondria (Fig. 5-7. Additionally, the decreased expression in several chloroplast-associated peptidyl-prolyl cis-trans isomerase paralogs, that participate in the assembly of protein complexes and redox regulation, strengthen the role of *Pp*HKT in maintaining ion homeostasis within the chloroplast for the well-functioning of this organelle.

The clearest phenotype in the *PpΔhkt* mutant is the deficiency in the production of sporophytes (Fig. 1), that could be related to an affectation in the physiology of the spermatozoid, as indicated by the diminished expression of several genes linked to the structure and movement of the flagellum, like dynein assembly factor 1, dynein heavy chain, intraflagellar transport proteins 81 and 46, and tubulin (Fig. 5E). The large down-regulation of gene Pp3c1_40600 with a -7.37-fold change) that encodes for an ADP ribosylation factor (*Pparl13b)*, that is involved in intraflagellar transport and cilia stability in humans (He et al., 2018), further indicate the possible malfunctioning of the spermatozoid as the factor responsible for the lack of sporophyte formation in the moss mutant (Meyberg et al., 2020). The production of few sporophytes in the *PpΔhkt* line when it was co-cultivated with the WT line, further indicate malfunctioning of the spermatozoid as the factor responsible for the lack of sporophyte production in the mutant line. Moreover, these assays also showed that *Pp*HKT could be participating in the early stages of sporophyte formation, as indicated by the deformed sporophyte (Fig. 1H), that could involve alterations in mitotic cell divisions to form the sporophyte capsule and/or meiotic cell divisions that occur during spore formation (Kofuji & Hasebe, 2014). These results indicate that both, flagellar structural and dynamic protein factors seem to be affected by the disruption of ion homeostasis within the chloroplast of the *PpΔhkt* mutant, and more importantly, considering that the spermatozoid possesses a chloroplast or plastid within its structure.

It is interesting that despite the supressed expression of many photosynthesis and chloroplast related genes, the mutant moss grew and developed a chlorotic gametophore which indicate that the upregulated genes help to sustain such development. From this information it seems that the coordinated upregulation of glycolysis, gluconeogenesis, β-oxidation/lipid metabolism, and respiration could explain the survival of the moss and larger size of the gametophores observed for the *PpΔhkt* mutant (Fig. 1). Within glycolysis SuSy, PPi-dependent phosphofructokinase (PFP), aldolase and triose phosphate isomerase are upregulated, activities that could lead to an increase in pyruvate, that together with the malate derived from the synchronised action of β-oxidation and the glyoxylate cycle, would be fed to gluconeogenesis for sugar synthesis. Additionally, if cytoplasmic glycolysis were stimulated, as indicated by the increased expression of sucrose synthase and PFP, the net yield of ATP would increase from 4 to 8, supplying the ATP required for all the anabolic processes involve in maintaining the development of the *PpΔhkt* mutant moss (Plaxton, 1996). Alternatively, acyl-CoA thioesterase and a fatty acid elongase complex are two upregulated transcripts that could lead to the synthesis of acetyl-CoA which is essential for production of cellular energy via the TCA and for synthesis of sugars and other carbon skeletons. The transcripts for phosphoglycerate kinase and aldolase, two enzymes within gluconeogenesis, would help to increase the synthesis of sugars to supply the TCA and the generation of cellular energy to assist in moss growth. An additional process that could function in a similar manner is the catabolism of the branched amino acids, leucine, isoleucine and valine, by the activity of Branched-chain Aminotransferases (BCAT; Pp3c14_21300 and Pp3c16_17790) that can serve as precursors of the TCA by synthesis of succinyl-CoA and acetyl-CoA, or as electron donors for the mitochondrial electron transport chain. In Arabidopsis, the *BCAT2* gene is expressed at a very low level, but transcript levels rise in response to various stresses like carbon starvation and hormone treatments (Matsui et al., 2008). Recently, the importance of respiration in *P. patens*, even when photosynthesis is active, has been postulated as the mechanism supplying cytosolic ATP, thus liberating its synthesis from photosynthesis regulation (Vera-Vives et al., 2024)/

Several genes encoding for transporters were upregulated, with two main groups corresponding to K^+^ and Zn^2+^ transporters, together with transporters for Fe^2+^, Cu^2+^ and Mn^2+^ (Fig. 8). These changes can be used to propose the increased need of the *PpΔhkt* mutant for ions required in maintaining the well-functioning of the organism, and particularly, of the chloroplast, where the ionic alterations generated by the absence of *Pp*HKT, very likely require the activity of the coordinated expression of several transporters to recover ion homeostasis and chloroplast functioning.

According to the likely localization of *Pp*HKT in the thylakoid membrane and its functioning as a Na^+^/K^+^ co-transporter, absence of this transporter would alter the thylakoid proton motive force (pmf), by preventing the dissipation of the membrane potential (Δψ), and leading to the establishment of a less acidic luminal pH (reduced ΔpH) that would affect the functioning of violaxanthin de-epoxidase (VDE; Pp3c3_13220) which requires an acidic pH for optimal functioning. This, in turn, would prevent the activation of non-photochemical quenching (NPQ) in the antenna of PSII, that would lead to a decrease in photosynthesis efficiency. This effect would be exacerbated in *PpΔhkt* where the expression of VDE suffered a two-fold decrease according to our RNAseq analysis (Table S2). Dissipation of Δψ as a mechanism to increase the contribution of ΔpH across the thylakoid membrane has been solved in other species through the activity of either, chloride channels (Herdean et al., 2016b) or K^+^/H^+^ exchangers (Armbruster et al., 2017; DeTar et al., 2021; Kunz et al., 2014b; C. Wang & Shikanai, 2019). Interestingly, in the *PpΔhkt* mutant transcriptome, the overexpression of two bestrophin-like chloride channels was observed, Pp3c1_34560 and Pp3c11_6680 (Fig. S5A), with a 1.4-fold increase, which could serve as alternative mechanisms for the dissipation of the thylakoid membrane potential in the absence of *Pp*HKT to ensure a partial recovery of chloroplast functioning and survival of the mutant. In *Arabidopsis thaliana*, two voltage-gated Cl^-^ channels (VCCN1 and VCCN2) have been identified in the thylakoid membrane; these channels belong to the bestrophin-like chloride channel family, which are conserved among prokaryotes and eukaryotes (Duan et al., 2016). According to structural and electrophysiological experiments, *At*VCCN1 and its orthologous in *Malus domestica*, *Md*VCCN1, function as anion selective channels to dissipate the thylakoid membrane potential and to sustain the ΔpH required for ATP synthesis (Duan et al., 2016; Hagino et al., 2022), supporting our interpretation. The transcriptome database from *P. patens* developmental stages (https://bar.utoronto.ca/efp_physcomitrella/cgi-bin/efpWeb.cgi) reveals that the moss bestrophin-like channel (Pp3c1_34560) and *Pp*HKT genes are highly expressed at the sporophyte S2 development (Fig. S5B); we confirmed these results by semi-quantitative RT-PCR from gametophores with early developing sporophytes was performed by a semi-quantitative RT-PCR (Fig. S5C), which suggest that *Pp*HKT and Pp3c1_34560 could function together to maintain the Δψ and ΔpH across the thylakoid membrane at this developmental stage. The participation of KEA’s or CLC’s as alternative mechanisms for the dissipation of thylakoid membrane potential can be ignored as none of the corresponding genes is significantly modified in *PpΔhkt.* In contrast to the strong phenotype observed for the *PpΔhkt* mutant (Fig. 1), single mutants of the thylakoid-localised K^+^/H^+^ exchanger KEA3, in Arabidopsis, did not cause a strong phenotype or alterations in photosynthesis activity (Kunz et al., 2014b). Mutation of the two other Arabidopsis KEA exchangers, KEA1 and KEA2 generated a dwarf phenotype with a diminished photosynthesis (Fv/Fm) ratio, although the location of these transporters has been assigned to the inner chloroplast envelop, where they should play a role in maintaining the stroma pH (Armbruster et al., 2014). In comparison to mutants affected in other chloroplast ion transporters, mutation of *PpHKT* has severe consequences to the moss phenotype, suggesting that control of the thylakoid pmf strongly depends on the activity of *Pp*HKT, and therefore, control of chloroplast ion homeostasis.

Finally, the central role of *Pp*HKT in the physiology of the moss chloroplast is reinforced by the strong modifications observed in the structure of the thylakoid in the *PpΔhkt* mutant (Fig. 3), that indicates that membrane formation, and possible protein complex arrangement depend on the establishment of a correct ion gradient, likely to be ΔpH-dependent, similarly to what has been observed for the *Atbest* mutant in Arabidopsis (Duan et al., 2016).

## CONCLUSION

The most contrasting feature of *Pp*HKT is its localization to the thylakoid where it must play a key role in controlling the pmf across the membrane, as a mechanism to activate NPQ for photoprotection. The central role of *Pp*HKT in the physiology of the moss is highlighted by the pleiotropic effects that are observed in the mutant, where the absence of the sporophyte, a diminished chlorophyll content and the larger size of the gametophores are among the most visible. Additional to these morphological changes, gene expression is modified significantly for close to one thousand genes, genetic changes that help to explain several of the phenotypes and serve to propose that glycolysis, respiration and lipid metabolism are few of the metabolic processes that are activated to help the mutant to almost complete its life cycle, with the exception of sporophyte formation which must be related to the malfunctioning of the spermatozoid. Association of *Pp*HKT to the chloroplast and functioning as a K^+^/Na^+^ co-transporter reflects the prokaryotic origin of these family of transporters.

## METHODOLOGY

### Plant material and growth conditions

*Physcomitrum patens* (‘‘Gransden’’ wild-type strain), the mutant *PpΔhkt1* (Haro et al., 2010)(Haro et al., 2010), and *Pp*HKT1-3xmNeonGreen transgenic moss lines were cultivated using standard conditions. The moss protonema tissue was collected, mechanically disrupted, and used to spread on sterile cellophane discs over Petri dishes with solid PpNH4 medium ([1.03 mM MgSO_4_, 1.86 mM KH_2_PO_4_, 3.3 mM Ca(NO_3_)_2_, 45 µM FeSO_4_, 2.72 mM (NH_4_)_2_-tartrate, 9.93 µM H_3_BO_4_, 220 nM CuSO_4_, 1.966 µM MnCl_2_, 231 nM CoCl_2_, 191 nM ZnSO_4_, 169 nM KI and 103 nM Na_2_MoO_4_], supplemented with 0.7% agar), and then incubated at 24°C, with a photoperiod of 16 h-light/8h-dark (70 µE m^-2^ s^-1^ of light) for 7 days.

### Protoplast isolation from *P. patens* protonema

Protoplasts were used as a starting material for morphological analysis and moss transformation following the protocol described below. For protoplast isolation, protonema from one plate of 7-day-old culture was digested with 1.5 ml of 2% driselase (Sigma-Aldrich) and 5 mL of 8.5% mannitol (Sigma-Aldrich) solution for 1 h incubating at room temperature in a shaker. Protoplasts were filtered through one layer of Miracloth (Merck-Millipore) to remove undigested tissue. The collected protoplasts were washed three times with 10 mL of 8.5% mannitol solution spinning at 250 *g* for 7 min in a 15 mL tube in a Sorvall ST8 centrifuge (Thermo Scientific) to finally be used for moss transformation or morphological experiments described in the next sections.

### Moss transformation

To obtain the *Pp*HKT-3xmNeonGreen transgenic moss line by the CRISPR-Cas9&HDR (homology-directed repair) system, the PEG protocol was employed as described (Mallett et al., 2019). Protoplasts were resuspended in a required volume of 3M buffer (9% mannitol, 150 mM MgCl_2_ 6H_2_O, and 0.1% MES, pH 5.6) to obtain a concentration of 2x10^6^ protoplasts mL^-1^. For protoplast transformation, an aliquot of 300 µl of protoplasts suspension was mixed with 15 µg of DNA, and 300 µL of PEG solution (40% PEG; 8.5% mannitol, 100 mM Ca(NO_3_)_2_·4H_2_O and 10 mM Tris-HCl, pH 8). The suspension was incubated for 10 min at room temperature and then heat shocked at 45°C for 3 min, followed by incubation for 10 min at room temperature. Then, 5 mL of 8.5% mannitol was added to the protoplast suspension and incubated at room temperature for 30 min. Finally, the protoplasts were recovered by centrifugation at 1600 rpm (250 *g*) for 5 min and resuspended in 0.5 mL of liquid plating medium (PpNH_4_ medium with 8.5% mannitol, 10 mM CaCl_2_), and plated on sterile cellophane disks over PRMB plates (PpNH_4_ medium supplemented with 6% mannitol, 10 mM CaCl_2_ and 0.7% agar); after four days, the cellophane was transferred to a PpNH4 plate supplemented with hygromycin (15 µg/mL). After one week, the cellophane was transferred to PpNH_4_ media without antibiotics, then, the cellophane was twice changed to a fresh solid PpNH4 medium every week. Finally, surviving plants were picked with sterile tweezers and inoculated onto solid PpNH4 medium without cellophane disk, allowing maximal growth by three weeks for genomic DNA extraction and transformants were verified by PCR.

### Morphological analyses

The analysis of gametophore morphology was started from protoplasts isolated from 7-day old protonema tissue resuspended in 0.5 μL liquid plating medium (PpNH_4_ medium supplemented with 8.5% Mannitol and 10 mM CaCl_2_) and spread on sterile cellophane disks over petri dishes with PRMB medium (PpNH_4_ medium supplemented with 6% Mannitol and 10 mM CaCl_2_) for four days, and then, transferred to PpNH_4_ medium for three days. Gametophore morphological analysis was performed in colonies from 7-day-old protonema tissue sections of 3x3 mm carefully placed with forceps on PpNH_4_ solid medium and grown under 24°C 16h-light/8h-dark photoperiod for four weeks. The gametophores with more than five phyillids and well-formed rhizoids (under stereomicroscope observation) were collected and counted. The moss colony and gametophores images were taken with a digital camera (Nikon 7500).

For reproductive structures and sporophyte analysis, 7-day old protonema tissue from regenerated protoplasts was propagated and the moss suspension was poured carefully over peat moss pellets (Jiffy-7, Jiffy Products International AS, Kristansand, Norway) in magenta boxes (Sigma-Aldrich) filled with ¾ parts of distilled water under sterile conditions. The moss was grown under 24°C on a 16h-light/8h-dark regime for four weeks until adult gametophores were observed; for induction of reproductive structures and sporophyte formation, the magenta boxes were transferred to a growth chamber at 16°C under a 16h-dark/8h-ligth short day photoperiod. Development of the reproductive structures, antheridia and archegonia, were monitored and carefully collected with sterile tweezers at 15, 21 and 28-day post induction (dpi). Sporophytes were observed and collected carefully with sterile tweezers under a stereoscopic microscope (Nikon Eclipse) after 28 dpi. The organs were observed under an inverted microscope (Nikon Eclipse), images were acquired with a digital camera (Nikon 7500) and adjusted bright and contrast to improve the shape of the reproductive structures.

### Cross fertilization between WT and *PpΔhkt* lines

To evaluate fertility of *PpΔhkt1* mutant line, a half of a peat pellet was inoculated with protonema from the WT line and the other half with protonema from the *PpΔhkt1* mutant. Gametophores were grown and exposed to gametangia-inducing conditions as described before. After four weeks, sporophytes were counted and analyzed.

### Chlorophyll Quantification

For chlorophyll extraction the protocol from Arnon (Arnon, 1949) was employed; gametophores from the *PpΔhkt1* mutant and WT lines were separately collected in microcentrifuge tubes and homogenized in 1 mL of 80% acetone; after incubation of the extract at room temperature for 2 h, samples were spined at 250 *g* in a microcentrifuge (Eppendorf), and the supernatant was collected in fresh tubes. Absorbance was measured at 645 and 663 nm using 80% acetone as blank in a spectrophotometer (Carry 60 UV-Vis, Agilent). Total chlorophyll content was calculated with the formula:

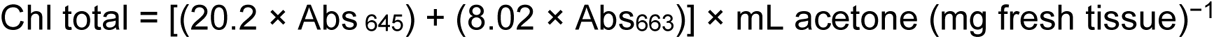

### Generation of 3XmNeon Knock-in line

The knock-in line HKT-3XmNeon line was generated by CRISPR-Cas9 & HDR (homology-directed repair) described by. The protospacer was designed using the CRISPOR online software (crispor.tefor.net, using *P. patens* (Phytozome V.11; (Concordet & Haeussler, 2018)) as the genome and *Streptococcus pyogenes* (5’NGGG3’) as the protospacer adjacent motif (PAM) parameters. One protospacer was chosen based on a high specificity score and a low off-target frequency, considering that the protospacer was closest to the stop codon of the *HKT* gene. The protospacer and its reverse complementary were synthesized as oligonucleotides (see TableS1) and annealed in a PCR reaction (500 pmol of each, 10 µL total volume, with the following setting conditions: 98°C 3 min, 0.1°C s^-1^ to oligo Tm, hold 10 min, 0.1°C s^-1^ to 25°C) to be joined by complementarity. The CCAT-sequence was added at the 5′end of each protospacer to generate a sticky end compatible with the BsaI-linearized pENTR-PpU6P-L1L2 entry vector. The protospacer was ligated to the entry vector using the Instant sticky-end ligation Master Mix (New England Biolabs) according with manufacturer’s specifications to obtain the pENTR-PpU6P-L1L2-hkt protospacer construct. The construct generated by ligation was recombined with pMH-Cas9-gate expression by an LR-clonase reaction following the manufacturer’s specifications.

To generate the homology arms for the tagging of *Pp*HKT, two 932 bp fragments upstream and downstream of the *PpHKT* initiation and stop codon, which correspond to the 5’ and 3’ arms, respectively, (Table S1) were used and amplified by PCR. The 5′ and 3′ arms were cloned by recombination by a BP-clonase reaction using the pDONR-P1P4 and pDONR-P3P2 entry vectors, respectively. The 5’ arm PAM sequence was modified by PCR site-directed mutagenesis (Wu et al., 2023), using the high fidelity Taq polymerase Q5 (New England Biolabs) according with manufacturer’s specifications and using the primers described in Table S1. The HDR construct to tag PpHKT at the C-terminus, was generated with the Multisite Gateway cloning system (Invitrogen), recombining the pDONR-B1-HKT-5′ arm-B4, pDONR-R4R3-3xmNeonGreen-C, pDONR-B3-HKT-3′ arm-B2 into the pGEM-gate destination vector by a triple LR reaction.

Finally, WT protoplasts from *P. patens* were co-transformed with the pMH-Cas9-HKT vector that contained the protospacer and the pGEM-HKT-3xmNeonGreen vector that contained the 3xmNeonGreen tag flanked by the 5′and 3′ PpHKT arms. The vectors used as backbones for the constructions were donated by the Bezanilla lab (https://www.addgene.org/Magdalena_Bezanilla/). Potential transgenic moss lines were analyzed by PCR using genomic DNA as template. Genomic DNA was purified using CTAB method described in (Yáñez-Domínguez et al., 2023). Positive lines amplified a 2634 bp fragment that include the 3’ end of *PpΔhkt1* gene and mNeonGreen gene. The primers used are described in Table S1.

### Fluorescence microscopy

To analyze the subcellular localization of the HKT-3XmNeon moss reporter line, pieces of tissue were placed onto microscope slides in water, covered with a cover slide, and observed by fluorescence microscopy using an inverted multiphotonic confocal microscope (Olympus FV1000), employing an excitation wavelength (λ_ex_) of 488 nm (0.5% power laser) and observed at λ_em_ 655 nm with a 60X oil immersion objective with a 1.3 NA. Images were processed and analyzed with ImageJ (Schindelin et al., 2012).

### Yeast complementation assay

To express *PpHKT* in yeast cells, the codifying sequence for *PpHKT* was amplified by PCR employing the primers described in Table S1. The PCR-amplified product was cloned into the Yep352-NHA1 vector (previously digested with PstI), by homologous recombination in *S. cerevisiae* BW31a cells, leaving the *PpHKT* ORF under the *NHA1* promoter, to yield the construction pYep352-PpHKT. The construction was confirmed by restriction analysis with PstI and by sequencing, using the oligonucleotides described in S-Table 1. The K^+^ uptake-deficient BYT12 (*MATa his3Δ leu2Δ met15Δ ura3Δ*; *trk1Δ::loxP trk2Δ::loxP)* strain and the Na^+^-sensitive BYT45 (*MATa his3Δ leu2Δ met15Δ ura3Δ*; *nha1D::loxP ena1–5D::loxP)* strain (Navarrete et al., 2010) of *S. cerevisiae* were transformed with the lithium acetate (LiAc) procedure. Yeast cells were grown at 30° C in standard rich medium [YPD; 1% yeast extract (Difco), 2% peptone (Difco) and 2% glucose (Sigma-Aldrich, Carlsbad, CA, USA)], or selective medium Yeast Nitrogen Base (YNB) containing 0.67% yeast nitrogen base without amino acids (Difco) and 2% glucose, supplemented with amino acids appropriated for auxotrophic growth. Amino acids were used at a concentration of 20 µg mL^-1^l (Sigma-Aldrich, Carlsbad, CA, USA). To determine the phenotype of the transformed cells, the drop-test assay was employed with 10-fold serial dilutions of the culture spotted on YNB medium supplemented with several concentrations of NaCl, KCl, or LiCl, cell growth was recorded after five days.

### RNA isolation and RNAseq analysis

After three weeks in gametangia induction conditions, one hundred gametophores were manually collected from the WT and *PpΔhkt* mutant lines grown on sterile peat pellets (Jiffy-7, Jiffy Products International AS, Kristansand, Norway) and ground in liquid nitrogen. RNA extraction was performed with TRIzol (Invitrogen) following manufacturer’s protocol. Three replicates of each sample (WT and *PpΔhkt1*) were processed and stored in Gen-Tegra RNA tubes (Gen-Tegra, Integen X). Library construction and RNA sequencing were carried out commercially by Novogen (Novogen Corporation Inc, USA). Bioinformatic data was processed by Novogen and iDEP application (Xijin Ge et al., 2020).

## Acknowledgments

This work was supported by Grant IN217423 from PAPIIT-UNAM to OP. We would like to thank Dr. Rosario Haro for the gift of the *PpΔhkt1* mutant and the *PpHKT* clone. CY-D and DL-G were supported by a Ph. D. scholarship from CONAHCYT-Mexico. CY-D, DL-G and DM-T received a Post Doc scholarship from the 2041 Fronteras-2019 CONAHCYT Grant to OP. KM-O received a Post Doc scholarship from DGAPA-UNAM. JC-G was awarded a Post Doc scholarship from CONAHCYT. We acknowledge the support from the LNMA and Dr. Veronica Rojo-Leon with the confocal microscopy.

## Author contributions

CY-D, KM-O, DL-G and DM-T carried out the experiments. KM-O, JC-G and OP analysed RNAseq data. All the authors participate in the writing of the article.

## Data availability statement

All relevant data can be found within the manuscript and its supporting materials.

## Supporting information

Additional Supporting Information may be found in the online version of this article.

## SUPPLEMENTARY FIGURE LEGENDS

**Figure S1. Deletion of *PpHKT* generates pleiotropic changes at the gametophyte and sporophyte stages. A** Mutation of *PpHKT* induces protonema branching that increases with age. Protonema from WT and *PpΔhkt* lines stained with Calcofluor White from 7 and 10 days (7-10-d) cultures. **B** Quantification of protonema area from WT and *PpΔhkt* lines. **C** Wild type (WT) moss colony grew on PpNH_4_ media formed normal round colonies (left panel) and *PpΔhkt* moss colony formed big gametophores and less protonema tissue (right panel), with the formation of gametophores after 30 d under a long day photoperiod condition.

**Figure S2. Deletion of *PpHKT* does not modify gonad development. Aa-c** Antheridia **Ad-f** archegonia from WT moss. **Ba-c** Antheridia **Bd-f** archegonia from *PpΔhkt* moss. Both lines were grown for four weeks under 16°C with a 16h-dark/8h-ligth photoperiod to induce gonad development. Similar observations were made in more than five moss cultures.

**FigureS3. Differential gene expression in the *Pp****Δ**hkt*** **mutant. A** Volcano plot showing the down- and up-regulated genes in *PpΔhkt* in comparison to the WT moss. **B** Number and percent of up- and down-regulated genes in *PpΔhkt.* **C** Heat map from two WT and two *PpΔhkt* samples showing four gene clusters associated with ion/solute transport (up-regulated), and photosynthesis, microtubule-based process, and cell projection organization (down-regulated) in *PpΔhkt*.

**Figure S4. Up-regulated genes in *PpΔhkt.* A** Molecular function GO groups induced by mutation of *Pp*HKT. **B** Up-regulated transcription factors in *PpΔhkt*.

**Figure S4. Bestrophin-like channels present in P. patens. A)** Phylogenetic tree of human bestrophin (HsBEST1, NP_004174.1), *Klebsiella pneumoniae* (KpBEST, WP_049046555.1) and bestrophin-like channels from plants like *Arabidopsis thaliana* AtVCCN1 (At3g61320.1), AtVCCN2 (At2g45870.1) and *Malus domestica* MdVCCN1 (XP_028961536.1), moss *Physcomitrium patens* PpBSTa-d (Pp3c3_22870V3.1.p, Pp3c4_15850V3.1.p, Pp3c1_34560V3.1.p and Pp3c11_6680V3.1.p), and algae *Clamydomonas reinhardtii* CreBST1-3 (Cre16.g662600_4532.1.p; Cre16.g663400_4532.1.p and Cre16.g663450_4532.1.p). Asterisks highlight the PpBTc24 and PpBSTd genes found in the *PpΔhkt* transcriptome **B)** Transcript expression of bestrophin-like ion channels and *PpHKT* along *P. patens* life cycle. **C)** Semi-quantitative PCR analysis for the expression of PpPBTc in WT (first row) and *PpΔhkt* mutant (second row), in comparison with *PpACT5,* a reference gene, in WT (third row) and *PpΔhkt* mutant (fourth row), and expression of *PpHKT* in WT (fifth row) and *PpΔhkt* mutant (sixth row). Samples were taken from gametophores with early developing sporophytes. Images shows PCR band amplification in a 1% agarose gel of the three genes evaluated at 23 to 35 cycles.

